# Dynamic regulation of CeA gene expression during acute and protracted abstinence from chronic binge drinking of male and female C57BL/6J mice

**DOI:** 10.1101/2024.02.02.578650

**Authors:** Hernán G. Méndez, Sofia Neira, Meghan E. Flanigan, Harold L. Haun, Kristen M. Boyt, Todd E. Thiele, Thomas L. Kash

## Abstract

Binge alcohol consumption is a major risk factor for developing Alcohol Use Disorder (AUD) and is associated with alcohol-related problems like accidental injury, acute alcohol poisoning, and black-outs. While there are numerous brain regions that have been shown to play a role in this AUD in humans and animal models, the central nucleus of the amygdala (CeA) has emerged as a critically important locus mediating binge alcohol consumption. In this study, we sought to understand how relative gene expression of key signaling molecules in the CeA changes during different periods of abstinence following bouts of binge drinking. To test this, we performed drinking in the dark (DID) on two separate cohorts of C57BL/6J mice and collected CeA brain tissue at one day (acute) and 7 days (protracted) abstinence after DID. We used qRTPCR to evaluate relative gene expression changes of 25 distinct genes of interest related to G protein-coupled receptors (GPCRs), neuropeptides, ion channel subunits, and enzymes that have been previously implicated in AUD. Our findings show that during acute abstinence CeA punches collected from female mice had upregulated relative mRNA expression of the gamma-aminobutyric acid receptor subunit alpha 2 (Gabra2), and the peptidase, angiotensinase c (Prcp). CeA punches from male mice at the same time point in abstinence had upregulated relative mRNA encoding for neuropeptide-related molecules, neuropeptide Y (Npy) and somatostatin (Sst), as well as the neuropeptide Y receptor Y2 (Npyr2) but downregulated, Glutamate ionotropic receptor NMDA type subunit 1 (Grin1). After protracted abstinence CeA punches collected from female mice had increased mRNA expression of corticotropin releasing hormone (Crh) and Npy. While CeA punches collected from male mice at the same timepoint had upregulated relative mRNA expression of Npy2r and downregulated mRNA expression of Gabra2, Grin1 and opioid receptor kappa 1 (Oprk1). Our findings support that there are differences in how the CeA of male and female respond to binge-alcohol exposure, highlighting the need to understand the implications of such differences in the context of AUD and binge drinking behavior.

## Introduction

Alcohol Use Disorder (AUD) affects individuals across the globe, and in the United States alone it has negatively impacted the lives of millions of individuals each year^1^. Sadly, in the United States alone around 140,000 people a year die from excessive alcohol use ^2^, and the yearly cost of excessive alcohol use ^3^ has by far exceeded the yearly cost of heart disease which is the leading cause of death in the United States ^4^. Surprisingly, contrary to popular belief, out of all drugs of abuse, alcohol has been shown to be the most harmful to the user and surrounding environment ^5^. This highlights the longstanding need to understand the underlying neurobiological drivers of excessive alcohol use and addiction.

Addiction has been classically described by the triphasic model which is characterized by periods of binge, withdrawal, and preoccupation/craving that ultimately may lead to relapse ^6^. Each period has been suggested to be driven by specific brain regions paired with a defined set of molecular key players^6^. We are particularly interested in the central nucleus of the amygdala (CeA), due to it being an important hub for anxiety and modulation of both binge alcohol consumption and withdrawal periods^7^. Furthermore, in the context of excessive alcohol consumption, binge drinking is the most predominant and financially burdensome and is highly associated with AUD^3,8^. Furthermore, the CeA has been shown to be responsible for controlling alcohol binge drinking and withdrawal^7,9–12^

In humans, the process of alcohol withdrawal is characterized by a period of abstinence and specific clinical manifestations that range from mild (ie: alcohol craving, anxiety, excessive sweating, headaches and etc.) to severe withdrawal (ie: seizures, delirium, etc.) ^13^. Notably, although withdrawal in mice does not show the same clinical manifestations that humans do, in preclinical models such as C57BL/6J mice, it has been shown that periods of abstinence after binge drinking are associated with various negative affective behaviors or withdrawal-like symptoms^13^. Most of these are heightened fear, and anxiety-like behaviors ^14^, which highlights the importance of the CeA, as the CeA is involved in many of these processes. Furthermore, similar to humans during withdrawal, these affective behaviors during abstinence can shift over time. It has been hypothesized that these negative affective states promote alcohol consumption via engagement of negative reinforcement^15^.

It is crucial to investigate which genes in the CeA are modulated during different periods of abstinence following binge drinking, as it has been proposed that there can be an emergence of a negative affective state over time^16^. Notably, most studies that explore induced changes in gene expression in the CeA, after a period of binge or chronic ethanol drinking, only focus on male rodents during acute periods of abstinence^11,12,17^. For these reasons, we decided to investigate how the relative expression of G protein-coupled receptors (GPCRs), neuropeptides, ion channel subunits, and enzymes, that have been associated with alcohol consumption are impacted by differing periods of abstinence, acute or protracted abstinence, following Drinking in the Dark (DID) in both male and female C57BL/6J.

## Methods

### Animals

Two independent cohorts of wild type C57BL/6J mice were obtained from the Jackson Laboratories (Stock #: 000664, Jackson Laboratories) for this study. All mice included in this study were 11 weeks of age upon initiation of experimentation. Each experiment included a water group that only received water for the duration of each. For experiment 1, the acute abstinence group, mice were subjected to 1 day of ethanol abstinence following DID. This group included male (n=10) and female (n=10) mice. The water group for experiment 1 included male (n=10) and female (n=10) mice. Alternatively, in experiment 2, the protracted abstinence group, mice were subjected to 7 days of ethanol abstinence following DID, as previously described^18^. Experiment 2 group, included male (n=10) and female (n=10) mice. The water group for experiment 2 included male (n=10) and female (n=10) mice. Mice were housed separately by sex in a 12:12h reverse dark-light cycle facility and had ad-libitum access to food and water. All mice were maintained in a pathogen-free facility at the University of North Carolina at Chapel Hill (UNC-CH) and experiments were conducted in accordance with the guidelines of UNC-CH’s Institutional Animal Care and Use Committee (IACUC). The study protocol was approved by the Administrative Panel on Laboratory Care at UNC-CH.

### Two Bottle Choice Drinking in the Dark

Mice were single-housed in a two-bottle choice cage, with ad-libitum Prolab Isopro RMH 3000 (LabDiet) and water bottles for habituation one week before experiment initiation. Following this habituation regimen, two-bottle choice Drinking in the Dark was performed in a reverse light cycle setting as in our lab’s previous published study ^18^, which consisted of 3 consecutive weeks 2hr access to alcohol and water (Monday-Wednesday) and 4-hour access to alcohol (Thursday).

On each drinking day, bottles of 20% ethanol v/v and water were pre-weighed and introduced to the cages 3 hours after the end of the light cycle. Mice subjected to DID each simultaneously received one ethanol bottle and one water bottle. Meanwhile, the water group for each cohort received two bottles of water. An empty cage was set up with water and alcohol bottles for experiment 2 to account for drip. All bottles were weighed after each session. The average weight of each mouse and the overall average drip of experiment 2 were used to calculate g\kg drinking for both experiment 1 and experiment 2.

### Tissue Collection

Following 1 day or 7 days of ethanol abstinence after DID, experiment 1 and 2 mice were anesthetized with isoflurane and immediately decapitated with their respective water groups. Brains were extracted and immediately placed in ice-cold 1X PBS buffer. Whole brains were placed on a cutting block and sectioned into slices, 1mm in thickness. Tissue from several brain regions was collected for use in additional studies, including a previous publication from our lab^18^. For the scope of this study, only tissue collected from the CeA was included. Brain punches from the CeA were bilaterally collected using a 1mm diameter tissue micro punch and immediately flash-frozen in a 1.5 mL tube on dry ice. Tissue was stored at –80°C until RNA isolation was conducted.

### RNA Isolation and Reverse Transcription Reaction

Samples were removed from the –80°C freezer and thawed on ice. CeA tissue was prepared for homogenization by placing one stainless steel bead (5mm) (Qiagen, PN69989) inside the tube. 300 uL volume of RLT Buffer from the Qiagen RNeasy Kit (Qiagen, PN74104) and 10 uL per mL of Beta-mercaptoethanol was added to each tube. The TissueLyser LT (Qiagen PN85600) was used to homogenize the samples for 4 minutes at 50 oscillations per second. Homogenate was then transferred to new tubes and RNA isolation was performed following the standard manufacturer’s protocol. RNA concentration was determined using a nanodrop machine (Thermofischer PN13400519). Reverse transcription of the RNA was performed to synthesize cDNA following the standard manufacturer’s instructions of the iScript Reverse Transcription Supermix for RT-qPCR kit (Bio-Rad PN 1708841). cDNA was submitted to the University of North Carolina Chapel Hill Advanced Analytics Core (UNCAAC) (Funding Code: P30 DK034987). The UNCAAC ran both the cDNA preamplification and the BioMark HD qRT-PCR of the 25 genes of interest (Table 1), and Actb was selected as the housekeeping gene (HKG).

**Table 1.**
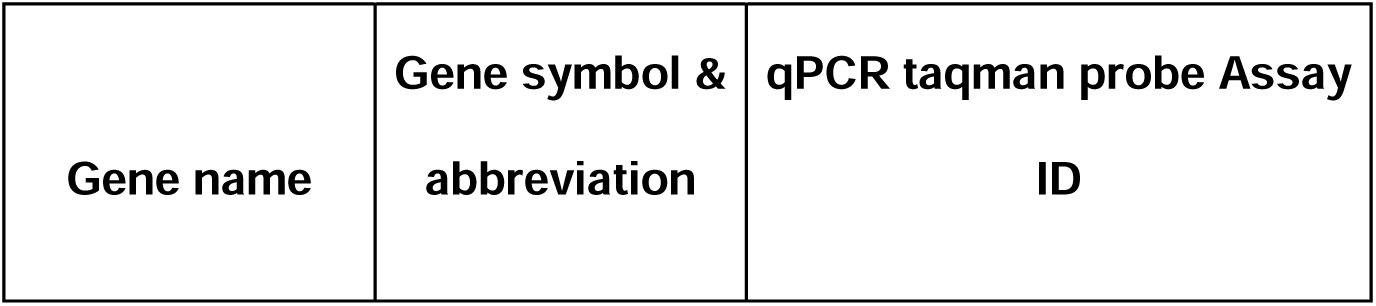
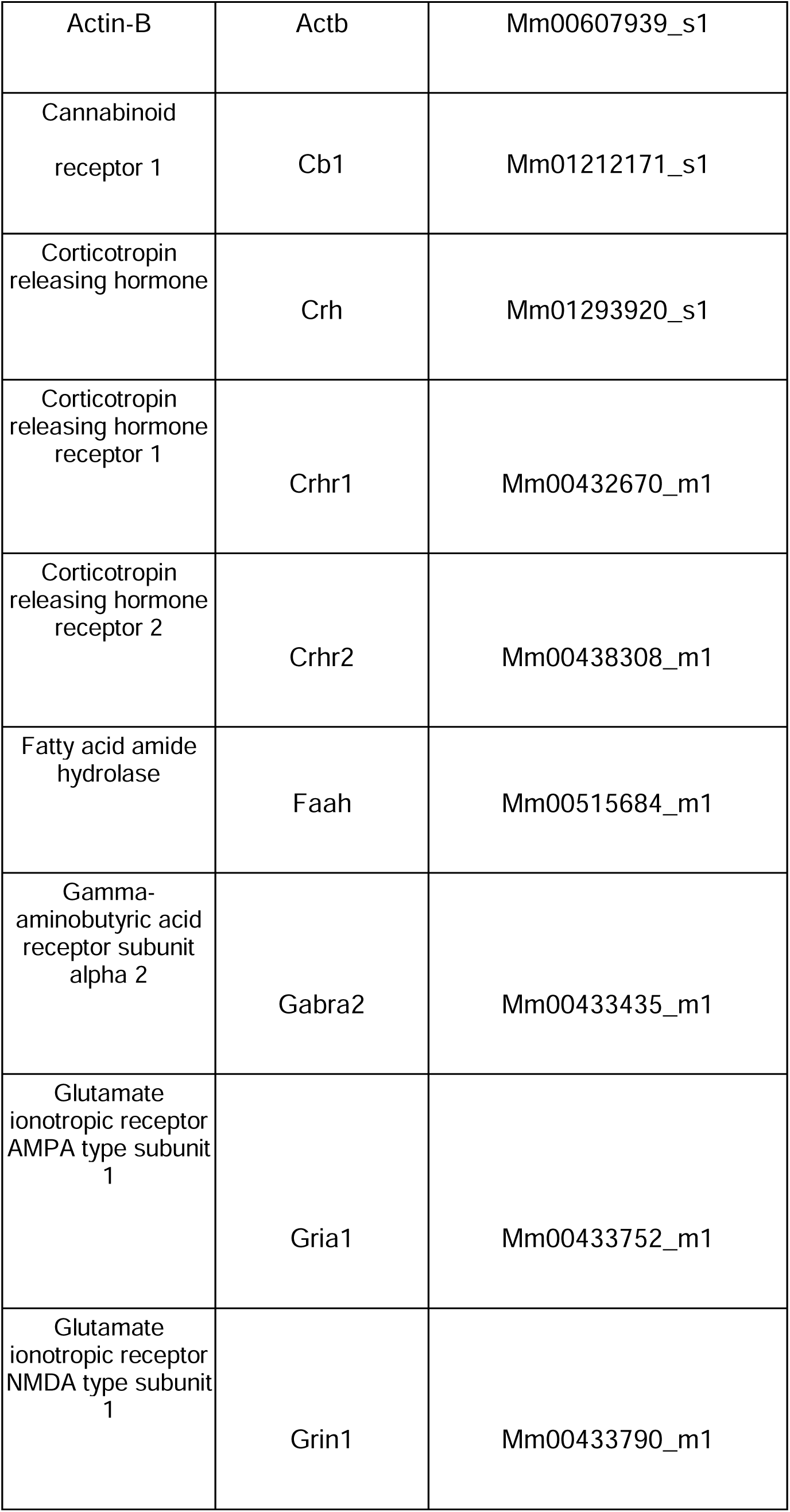
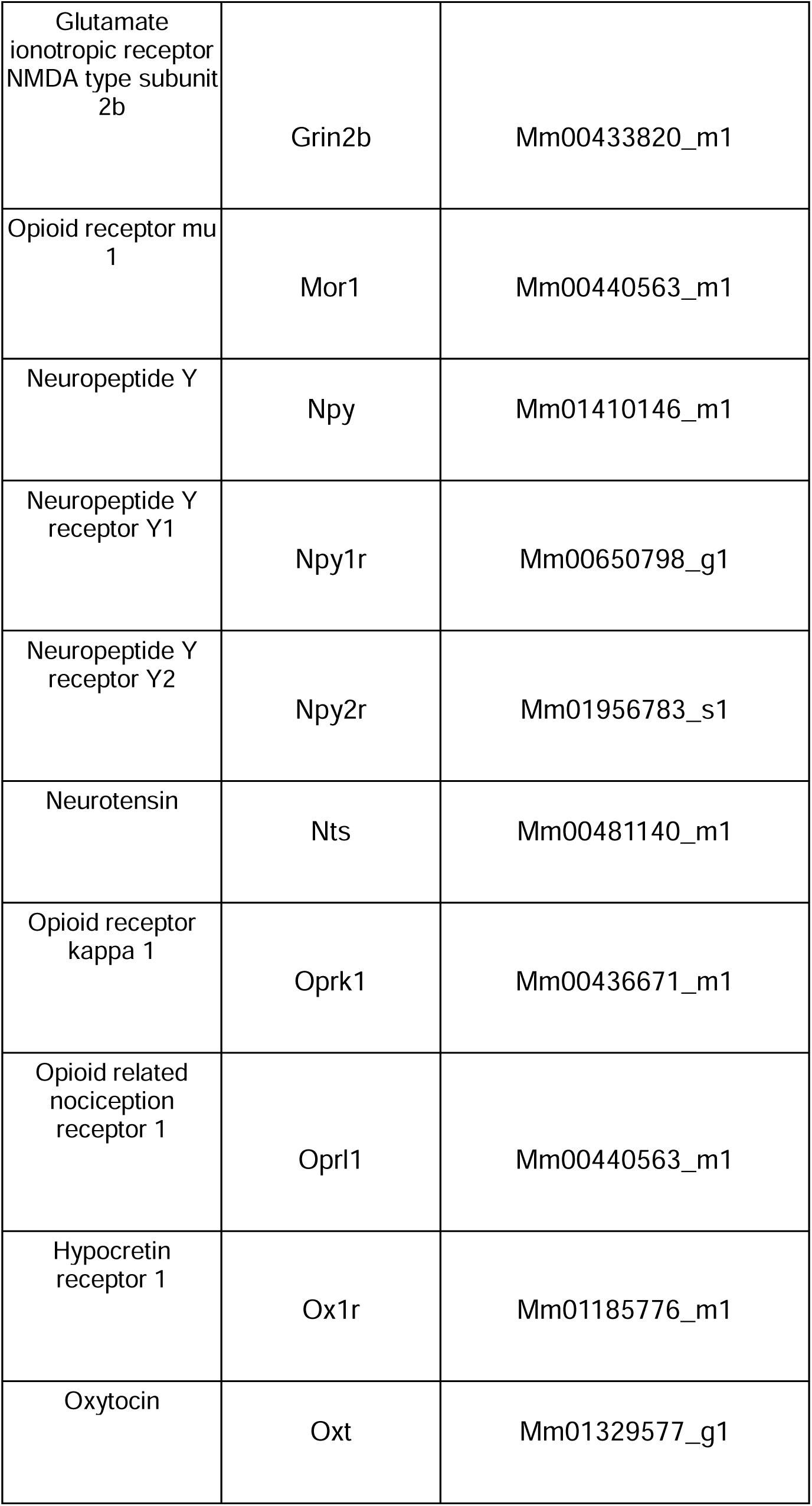
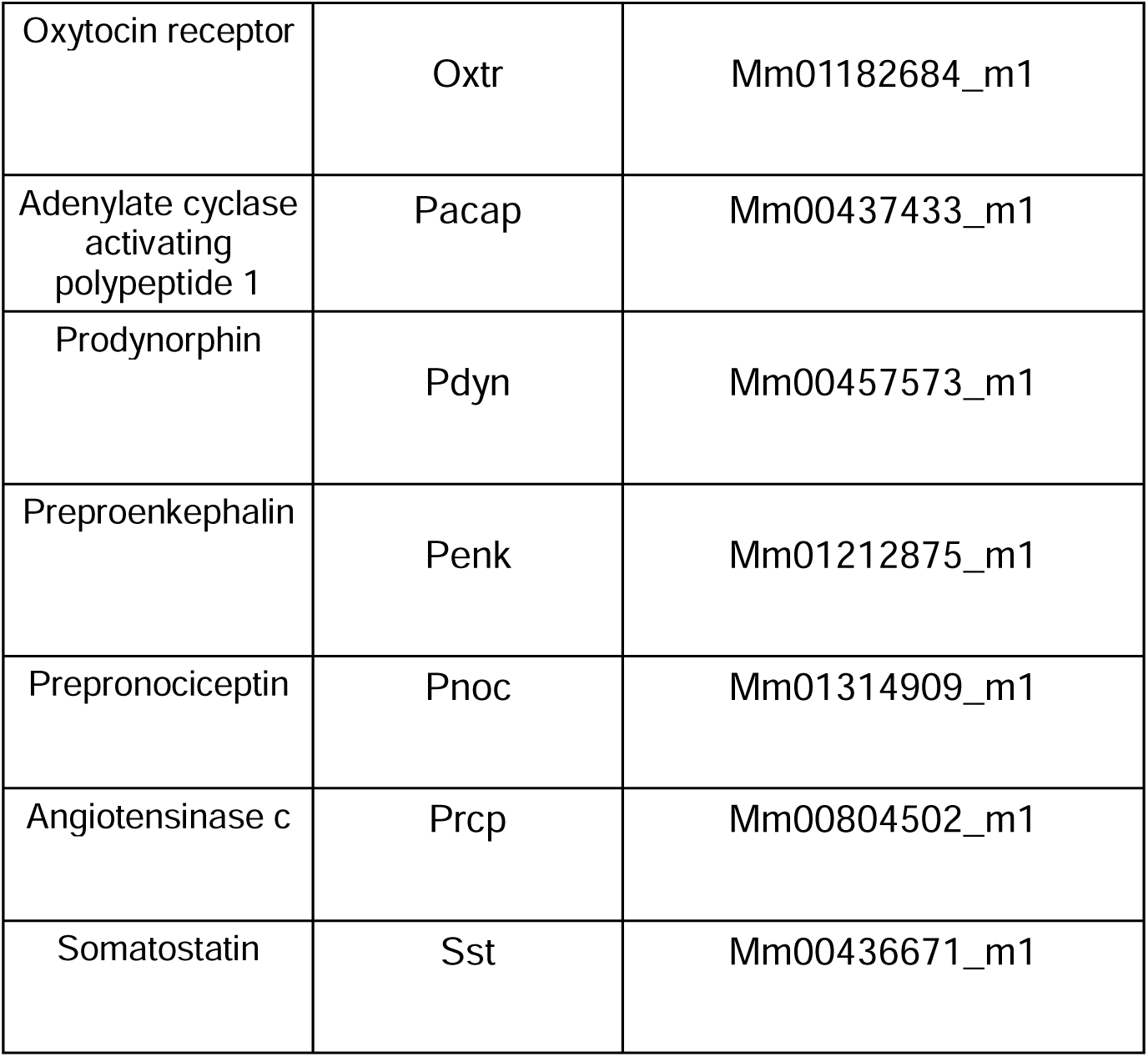
Taqman Probe & Assay ID (Thermos Fisher Scientific)

### cDNA Preamplification

Specific target amplification (STA) was performed according to manufacturer’s instructions (Standard BioTools PN 100-5876, https://tinyurl.com/yc4hxw2d). Thermal cycling was performed with the following settings: 2 min at 95 °C, 18 cycles of 15 sec at 95 °C and 4 min at 60 °C, held at 4 °C (C1000 Touch Thermal Cycler; Bio-Rad). Preamplified cDNA was immediately diluted 1:5 with 20 uL DNA Suspension Buffer and stored at –20 °C until use.

### BioMark HD qRT-PCR

Gene expression qRT-PCR was performed according to manufacturer’s instructions (Standard BioTools PN 100-2638, https://tinyurl.com/bddb5fyt), using a 96.96 Dynamic Array IFC (Standard BioTools PN BMK-M-96.96) following the manufacturer-provided layout. The TaqMan assay mixes and cDNA sample pre-mixes were loaded onto the IFC using the HX Loader (Standard BioTools). qPCR was performed using the BioMark HD instrument (Standard BioTools) with the “GE 96×96 Fast v1” thermal protocol.

### qRT-PCR Data Analysis

Since female mice consumed significantly more ethanol than male mice cumulatively, male and female groups within each experimental group were analyzed separately. 4 technical replicates were used to calculate relative gene expression. Relative gene expression was calculated using the 2^-(ΔΔCT) method, relative to male or female water group ActinB expression. Experimental groups 1 and 2 were normalized to their respective male or female water group.

## Statistical analysis

For statistical analysis purposes, male and female groups were analyzed separately because female mice drank significantly more ethanol than male mice. GraphPad Prism software version 10.0.2 was used for all statistical analyses. A repeated measure 2-way ANOVA and mixed effect analysis was performed to analyze differences in weekly ethanol intake. Šídák multiple comparison post hoc tests were performed whenever a main interaction was significant. Unpaired two-tailed student’s t-test or Mann-Whitney test was performed to assess differences in cumulative ethanol intake and relative gene expression. Pearson correlation matrix was performed to assess for correlations between gene expression changes, to correct for unequal N, we used a pairwise deletion method. A p-value less than 0.05 is considered significant. All data are plotted with Mean +-Standard error of the mean (SEM).

## Data Exclusion

The Grubbs test was performed to identify outliers in the data; which were excluded from the analysis. Also, some samples were excluded due to accidental loss during the RNA isolation period.

## Results

To study the impact of acute and protracted periods of ethanol abstinence after DID on relative gene expression in the CeA of C57BL/6J mice, we needed to first understand the alcohol binge-drinking behavior of our mice as they underwent the DID regime **(Figure 1A)**. To do this, we conducted two separate experiments that used a modified two-bottle choice DID regime, that was composed of a period of 3 consecutive weeks of 3 days of 2-hour drinking intervals and a fourth day of a 4-hour drinking interval. We found that both weekly and cumulative alcohol intake increased to binge levels. Notably, female mice drank significantly more alcohol than male mice in the third week of DID and the cumulatively across the three weeks in the acute abstinence group **(Figure 1B)** Repeated measures two-way ANOVA, Sex F(1,18)=13.07, P-value=0.002, Week F (1.984, 35.71) = 9.180, P-value < 0.001, Week and Sex interaction F (2, 36) = 0.8740, P-value=0.426, Šídák’s multiple comparisons test, P-value=0.005, Figure1C, Unpaired Student T-test: t=2.291, P-value=0.0342). Female mice also consumed more alcohol than males in the protracted abstinence group (**Figure 1D**) Mixed effect analysis, Sex F (1, 18) = 7.710, P-value=0.012, Week F (1.736, 30.39) = 2.101, P-value=0.145, Week and Sex interaction F (2, 35) = 1.927, P-value=0.161, Šídák’s multiple comparisons test, **(Figure 1E)**, P-value=0.020, Unpaired Student T-test: t=2.594, P-value=0.0189). Due to these differences in alcohol intake, we separated male and female analysis for the rest of this study.

**Figure 1:**
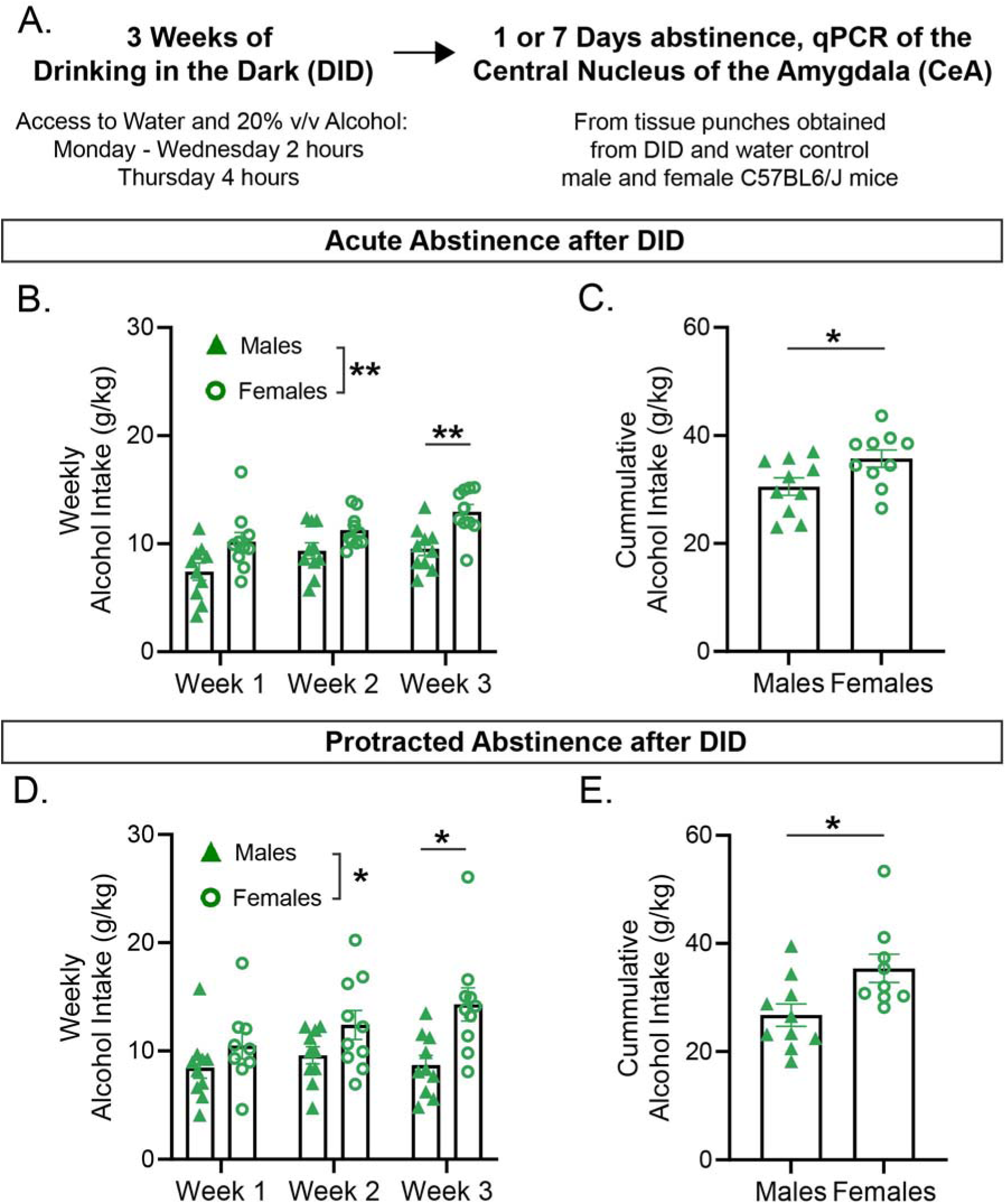
Drinking in the Dark Alcohol Intake. **A**) Schematic highlighting the experimental set up where tissue punches from the CeA of C57BL6/J mice were obtained at acute abstinence (1 day of abstinence following DID) or 7 protracted abstinence (7 days of abstinence following DID). **B)** Weekly alcohol intake of both male and female mice that underwent 3 cycles of DID and acute abstinence (1 day of abstinence following DID), male (n=10), female (n=10). **C)** Cumulative alcohol intake g/kg during the 3 cycles of DID of male (n=10), and female (n=10), mice that underwent 3 cycles of DID and acute abstinence (1 day of abstinence following DID). **D)** Weekly alcohol intake of both male and female mice that underwent 3 cycles of DID and protracted abstinence (7 days of abstinence following DID), (male n=10, females n=9-10). **E)** Cumulative alcohol intake g/kg during the 3 cycles of DID of male (n=10) and female (n=9) mice that underwent 3 cycles of DID and protracted abstinence (7 days of abstinence following DID). Male are represented as filled green triangles; female are represented as open green circles. Error bars are depicted as ±SEM. The following symbols represent, *= p-value < 0.05, **= p-value < 0.01, depicting statistically significance.

We then wanted to inquire how differing periods of withdrawal after DID affect relative gene expression of our 25 genes of interest in the CeA. The 25 genes consisted of GPRCRs, neuropeptides, subunits of ion channels and enzymes of interest in the CeA of male or female mice. To do this we extracted mRNA from the CeA from male or female mice that underwent either acute or protracted alcohol abstinence after DID and water group, and using qRTPCR we probed for relative gene expression changes. Interestingly, the CeA of male mice only showed significant gene expression level changes in 4 out of 25 genes of interest for both periods of abstinence. Specifically, male mice that went through acute abstinence after DID displayed 3 significantly upregulated genes(**Figure 2 A-C)**, Npy (Unpaired Student T-test: t=2.615, P-value=0.0176), Npy2r (Unpaired Student T-test: t=3.827, P-value=0.0015), and Sst (Unpaired Student T-test: t=2.646, P-value=0.0170). In addition, we observed 1 significantly downregulated gene during acute abstinence in males (**Figure 2D)**, Grin1 (Unpaired Student T-test: t=4.417, P-value=0.0004). In male mice that underwent a period of protracted abstinence after DID we observed 1 significantly upregulated gene: (**Figure 2F**), Npy2r (Unpaired Student T-test: t=2.692, P-value=0.0154), and 3 significantly downregulated genes (**Figure 2E, G-H)**, Gabra2 (Mann-Whitney test: U=15, P-value=0.0249), Grin1 (Unpaired Student T-test: t=2.653, P-value=0.0174), and, Oprk1 (Unpaired Student T-test: t=2.121, P-value=0.0499) compared to their water group. These results suggest that Npy2r is a common gene that is upregulated during periods of abstinence from binge drinking in males, regardless of whether this abstinence was acute or protracted.

**Figure 2:**
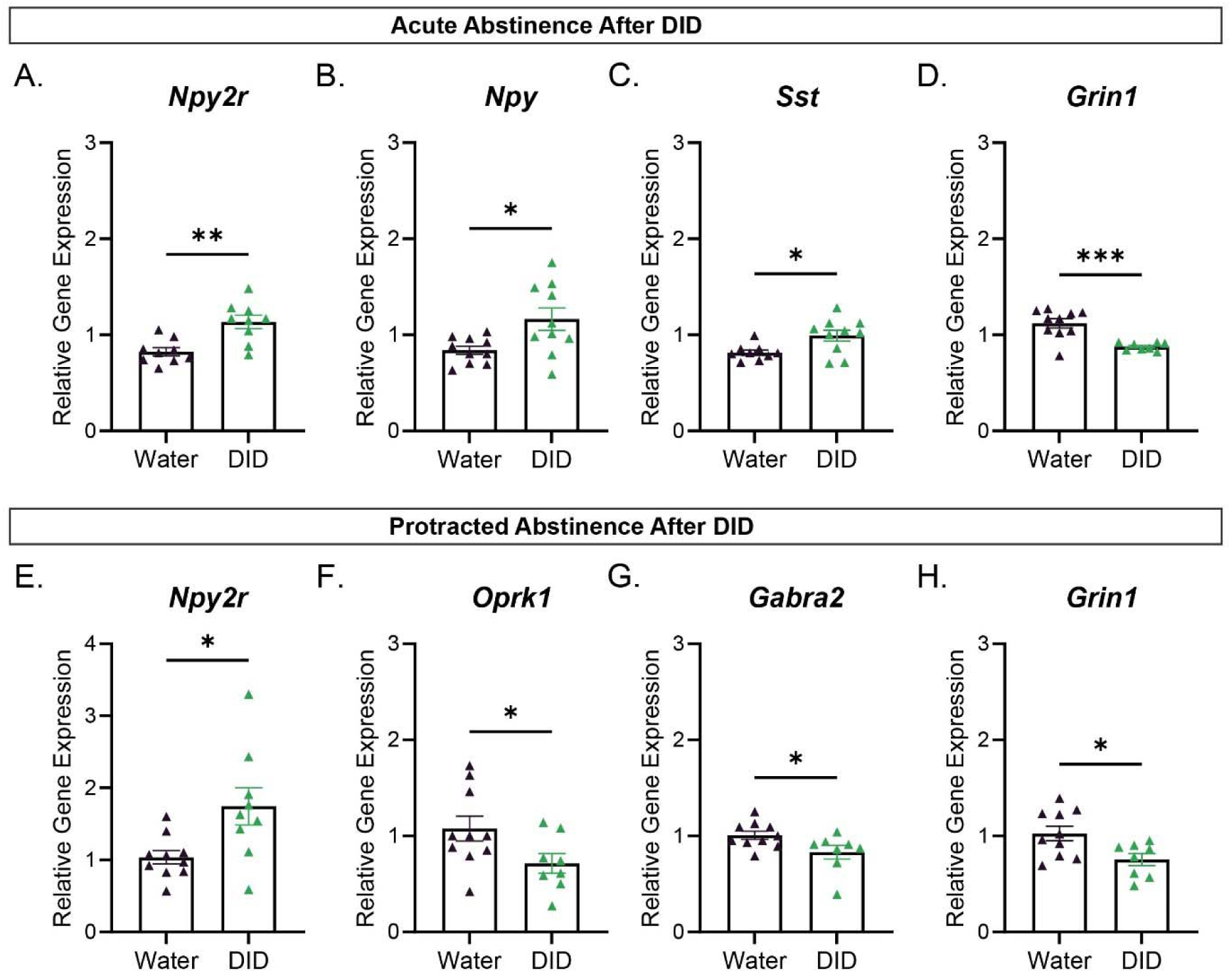
Male relative gene expression relative to housekeeping gene Actin B via qPCR. (**A-D**) Statistically significant relative gene expression levels of CeA derived genes from mice that underwent 1 day of abstinence following DID compared to the respective water group. **(E-H)** Statistically significant relative gene expression levels of CeA derived genes from mice that underwent protracted abstinence (7 days of abstinence following DID) compare to the water group. Error bars are depicted as ±SEM. The following symbols represent, *= p-value < 0.05, **= p-value < 0.01, depicting statistically significance. Water group n=9-10, DID group n=9-10.

Interestingly, the CeA of female mice only showed significant gene expression level changes in 2 out of 25 genes of interest for both periods of withdrawal. Specifically, female mice that experienced acute abstinence after DID had 2 significantly upregulated genes, (**Figure 3 A-B**), Gabra2 (Mann-Whitney test: U=11.50, P-value=0.0045) and Prcp (Unpaired Student T-test: t=2.207, P-value=0.0413). There were no significantly downregulated genes at this time point compared to the female water group. (**Figure 3C-D**), Furthermore, female mice that went through a period of protracted abstinence after DID did not display any significantly upregulated genes, but did display 2 significantly downregulated genes: Crh (Unpaired Student T-test: t=2.332, P-value=0.0322) and Npy (Mann-Whitney test: U=16, P-value=0.0167) compared to the female water group.

**Figure 3:**
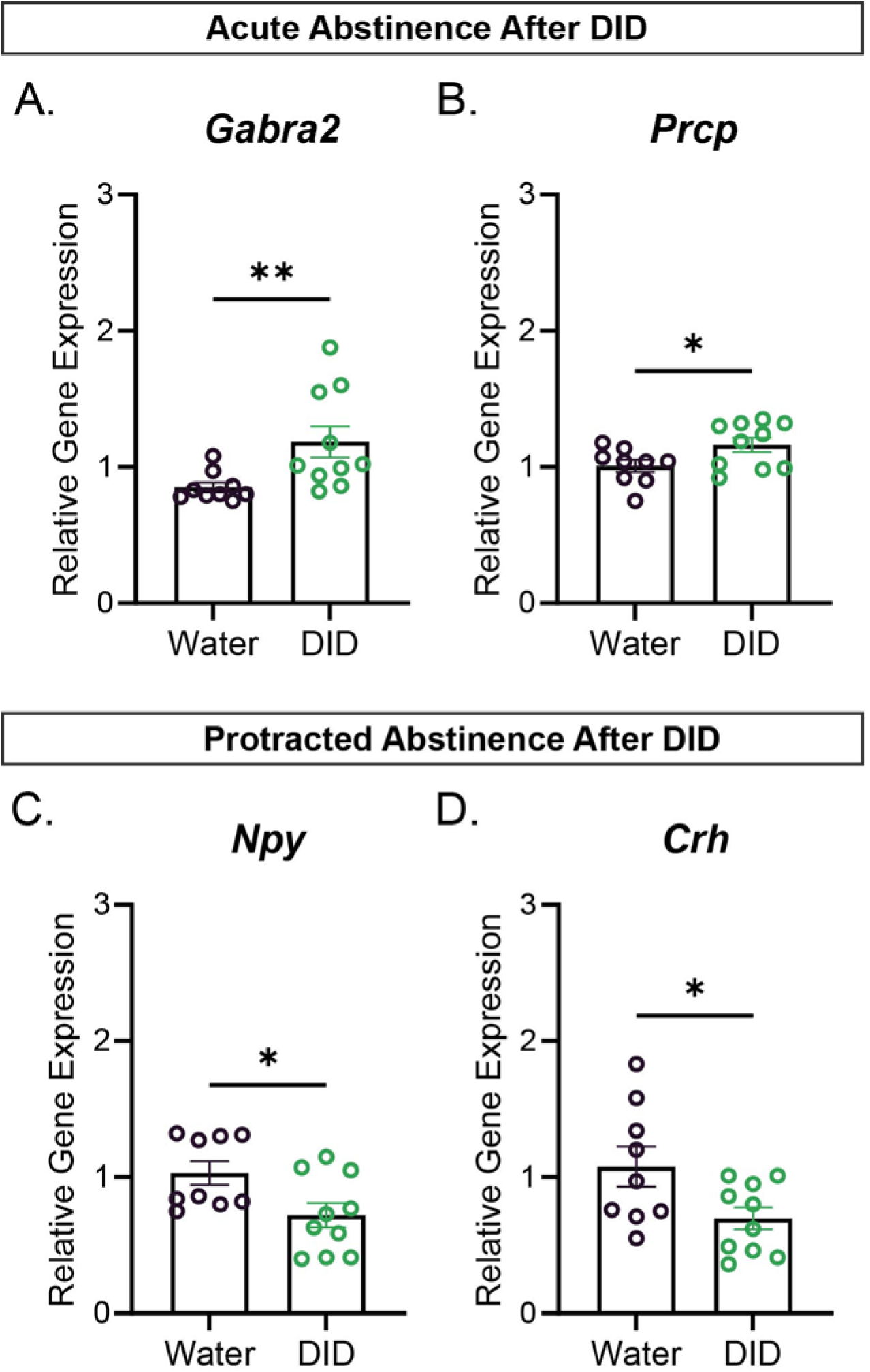
Female relative gene expression relative to housekeeping gene Actin B via qPCR: **(A-B)** Statistically significant relative gene expression levels of CeA derived genes from mice that underwent acute abstinence (1 day of abstinence after DID) compared to the water group. **(C-D)** Statistically significant relative gene expression levels of CeA derived genes from mice that underwent protracted abstinence (7 days of abstinence after DID) compared to the water group. Error bars are depicted as ±SEM. The following symbols represent, *= p-value < 0.05, **= p-value < 0.01, depicting statistically significance. Water group n=9-10, DID group n=9-10.

Next, we wanted to investigate if differing periods of abstinence after DID would affect the gene expression dynamics of our 25 gene of interest in the CeA of both male and female mice. To do this we used the relative gene expression data and a Pearson correlation analysis. We found a number of genes whose expression levels were significantly positively correlated across both male water group (**Figure 4A and 4C).** These positively correlated groups included **(Figure 4B**), Cb1/ Gabra2/Gria1/Grin1, Npy/Gria1, Pdyn/Penk, and Gria1/Faah/Grin1. Negatively correlated genes included Cb1/Oxt and Oxt/Grin2b. However, when mice were exposed to DID and given a day of abstinence, only the negative correlation between Cb1/Oxt was conserved **(Figure 4D).** Interestingly, for the male protracted abstinence after DID group only Cb1/Gria1 and Pdyn/Penk maintained their positive correlation, suggesting a change in coordinated gene expression.

**Figure 4:**
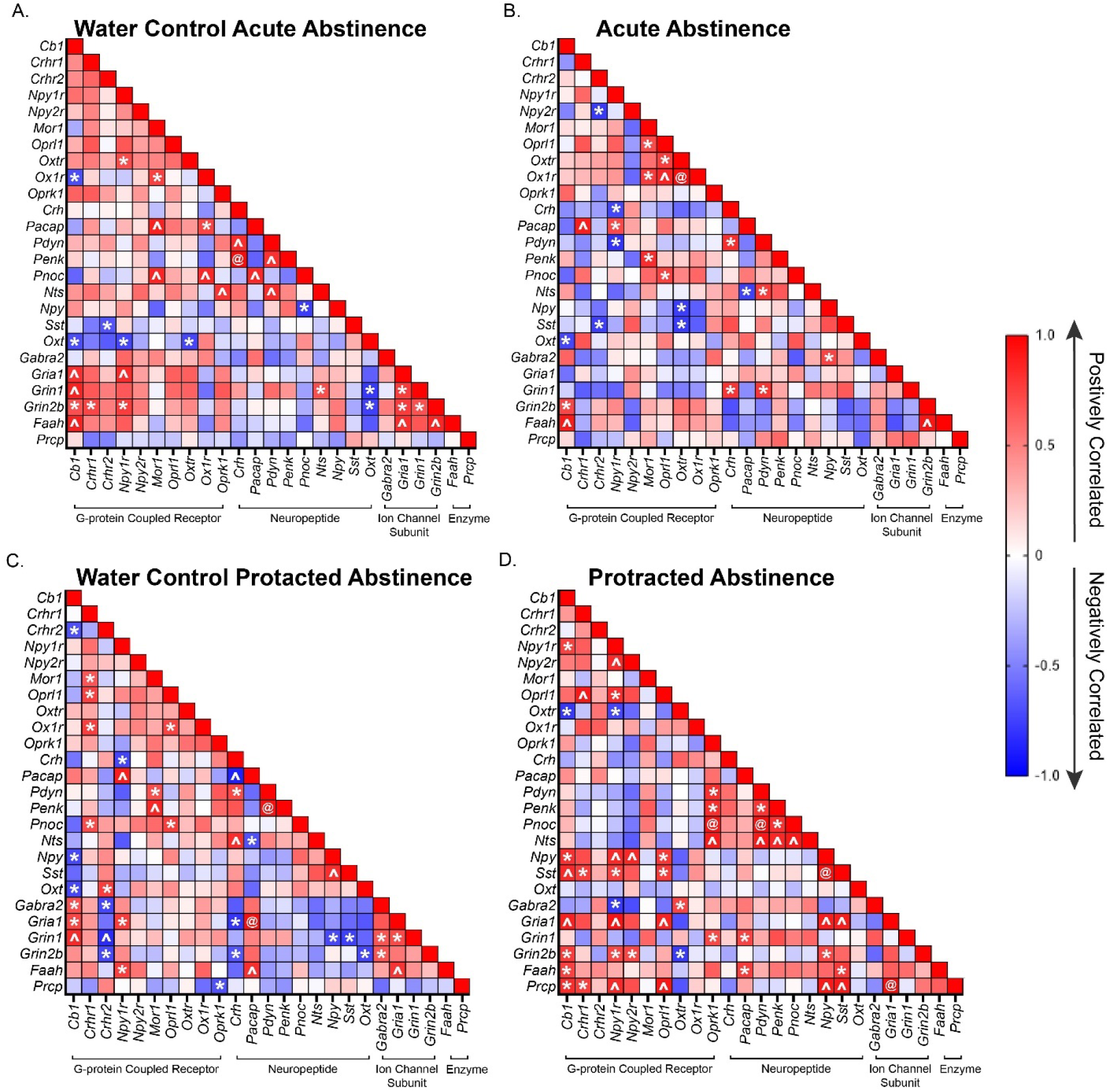
Male Pearson correlation of relative gene expression. **A**) Pearson correlation matrix of relative gene expression of genes of interest in the CeA of male mice that only received water. The CeA was isolated at the same time as the acute abstinence group (1 day of abstinence after DID after DID). **B)** Pearson correlation matrix of relative gene expression of CeA genes of male mice that underwent acute abstinence (1 day of abstinence after 3 cycles of DID). **C)** Pearson correlation matrix of relative gene expression of CeA genes of interest of male mice that only received water. The CeA of these was isolated at the same time as the protracted abstinence group (7 days of abstinence after DID). **D)** Pearson correlation matrix of relative gene expression of CeA genes of male mice that underwent protracted abstinence (7 days of abstinence after 3 cycles of DID). The following symbols represent, *= p-value < 0.05, ^= p-value < 0.01, @= p-value < 0.001, depicting statistically significant correlation.

In female mice, the Pearson correlation analysis showed a number of genes that were significantly positively correlated across both of the female water group **(Figure 5A and 5C)**, Crhr2/Pnoc/Pacap, Ox1r/Pacap, Crh /Penk, and Pacap/Faah. Notably, after DID **(Figure 4B and 4D)**, the only conserved relationship was the strong positive correlation between Pacap/Faah in the protracted abstinence group while none were conserved for in the acute abstinence group, suggesting a sex difference in coordinated regulation of these important alcohol consumption related genes in the CeA.

**Figure 5:**
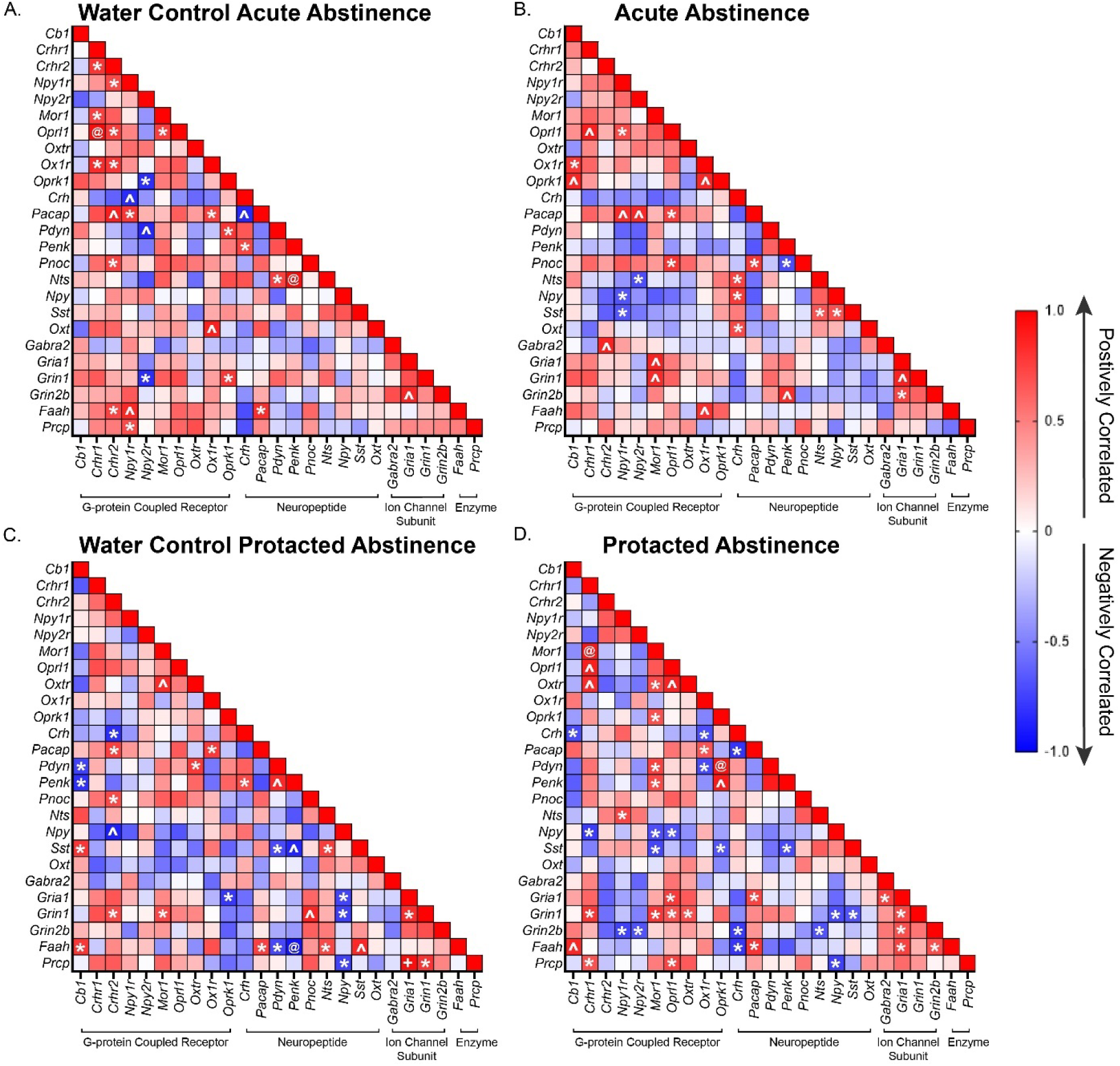
Female Pearson correlation of relative gene expression. A) Pearson correlation matrix of relative gene expression of genes of interest in the CeA of female mice that only received water. The CeA was isolated at the same time as the acute abstinence group (1 day of abstinence after DID after DID). **B)** Pearson correlation matrix of relative gene expression of CeA genes of female mice that underwent acute abstinence (1 day of abstinence after 3 cycles of DID). **C)** Pearson correlation matrix of relative gene expression of CeA genes of interest of female mice that only received water. The CeA of these was isolated at the same time as the protracted abstinence group (7 days of abstinence after DID). **D)** Pearson correlation matrix of relative gene expression of CeA genes of female mice that underwent protracted abstinence (7 days of abstinence after 3 cycles of DID). The following symbols represent, *= p-value < 0.05, ^= p-value < 0.01, @= p-value < 0.001, depicting statistically significant correlation.

## Discussion

Our goal was to investigate differences in relative gene expressions in the CeA at 1 or 7 days of abstinence from repeated cycles of DID in both female and male mice. This is important because the CeA is an integral control hub for binge drinking and withdrawal, but not much is known about the changing landscape of the genes during different periods of abstinence after binge drinking, much less about how it varies in males or female mice. We focused our study on genes that are heavily associated with alcohol related phenotypes.

In male mice, we found opposing effects of alcohol on the Npy system when compared to the female group. Notably, in the CeA of male mice that underwent 1 day of abstinence after DID, neuropeptide and receptor Npy and Npy2r were upregulated **(Figure 2 A-B)**. As previously discussed, CeA Npy decreases binge drinking in mice, although this same group shows a reduction in Npy protein following 24 hrs of abstinence after DID in males. This study found a similar change in Y2R mRNA level 24 hrs after DID, as well as an increase in Npy2r protein presence. ^19^ Therefore, our study is consistent with previous reports showing increased Npy2r expression in males following alcohol exposure, and highlight the potentially important role that Npy2r may be playing in excessive alcohol consumption, specifically via the CeA.

Sst was also upregulated in the acute abstinence after DID male group **(Figure 2C)**, but immunoreactivity data in the literature suggests that Sst is downregulated after DID in the CeA. ^20^. These differences in results could be due to differences in timing of when the tissue was collected and that we are only measuring mRNA levels. Interestingly, they showed that intra injection of Sst into the CeA reduces binge drinking and anxiety-like behavior. ^20^.

Males at the protracted abstinence after DID timepoint had downregulated Gabra2, Grin1 and Oprk1 **(Figure 2F, H-I)**. Consistent with our finding of downregulated Gabra2 in male mice, a study found downregulation of Gabra2 in postmortem CeA tissue of human males with a history of AUD^21^. We were surprised to see a reduction on Oprk1 expression, as previous work from our lab and others found that CeA Oprk1 deletion or blocking via pharmacological antagonist, reduces binge drinking, albeit only in male mice. ^22,23^ However, a study from the Zweifel lab found that genetic deletion of Oprk1 increases anxiety-like behavior in mice, suggesting a potential complex anxiety/alcohol intake interaction. ^24^ In addition, another study found that Oprk1 levels are reduced in the amygdala of humans with AUD. While this report did not specifically study the CeA, it highlights the potential involvement of Oprk1 in alcohol related behaviors across multiple species. ^25^

Grin1 was downregulated across both abstinence periods after DID in male mice. Grin1 is known to be an NMDA receptor ion channel subunit. While the NMDAR has long been known to be a target of alcohol, relatively little work has been done examining functional changes in subunit expression during binge drinking and subsequent withdrawal periods. Preclinical transgenic mice studies not isolated to the CeA where they used the mouse model Grin1 D481N, which has low binding affinity for glycine in the R1 subunit, showed that Grin1 D481N mice that were exposed to alcohol had fewer signs of anxiety-like behaviors compared to wild-type mice ^26^. On the other hand, another group that used cell culture differentiated cortical neurons from human embryonic stem cells found that after 7 days of ethanol treatment and 1 day of no ethanol, Grin1 was upregulated via qRTPCR measurements. ^27^ Another study using cells derived from AUD individuals found that Grin1 relative gene expression was also upregulated after 7 days of alcohol treatment. ^28^ Due to the lack of research about how binge drinking affects Grin1 dynamics, more research needs to be done regarding its role during binge drinking and during periods of withdrawal after binge drinking, especially in the context of the CeA.

In the CeA of the female acute abstinence group after DID only Gabra2 and Prcp were upregulated **(Figure 3 A-B**); however, this did not extend to the protracted abstinence time point **(Supplementary** Figure 3), suggesting that these genes are modulated by recent alcohol intake but not protracted abstinence. Gabra2 is a subunit of the GABA receptor, and mutations to the Gabra2 gene in humans have been associated with an increasing risk of suffering from AUD ^21^. Furthermore, in human female subjects that suffered from AUD with cirrhosis, higher levels of Gabra2 were found in the cortical region ^29^. Furthermore, a study in alcohol-preferring rats found that CeA Gabra2 modulates binge drinking behavior via Toll-like receptor 4 activation. ^30^ Therefore, our results are consistent with previous data in females and suggest that alcohol-induced changes in Gabra2 expression during abstinence may promote future binge drinking.

Prcp is known to regulate food consumption due to changes in the paraventricular nucleus VN of the hypothalamus (PVN), although these studies have not included female mice in their research. ^31^ In our studies, we found this enzyme was upregulated during acute abstinence after DID in female mice, **(Figure 5A)** which is a novel finding that needs to be further addressed in the field. This is particularly interesting as this is a peptidase that can degrade the neuropeptide, Alpha-MSH^32^. Alpha MSH can drive increased anxiety-like behavior via activity at the melanocortin 4 receptor, and studies have found reductions of alpha-MSH in the CeA following binge-like alcohol exposure^33^. It is possible that local actions of this enzyme may play an important role in this process, and this should be tested in future studies.

Crh and Npy were downregulated in the CeA of female mice after protracted abstinence after DID **(Figure 3 C-D)**. Crh in the CeA has been a target of many studies that investigate alcohol’s influence in the CeA. It was surprising to see that Crh was downregulated in the protracted period of withdrawal, since most of the immunohistochemistry data in the field shows increased levels of the peptide hormone corticotropin release factor (CRF) during withdrawal periods after DID. However, we cannot discount the possibility of changing protein levels since our study focused on the mRNA level. This down-regulation of Crh mRNA may reflect a compensatory change that counteracts this increase in Crh peptide. Npy is implicated in protection against drinking and anxiety-like behavior, especially during periods of withdrawal, for instance, Npy infusion in to the lateral ventricle attenuates alcohol consumption, potentially via actions in the CeA^19^. Notably, many of the studies focusing on the actions of these peptides have been performed in male mice. We speculate that this reduction in NPY may promote a heightened state of anxiety, as NPY is robustly anxiolytic.

There are some limitations of our study. One key limitation is that female and male mice drank different quantities of alcohol, therefore we were not able to control for differences in alcohol dosage to make a direct comparison between sexes. It is possible that these differences in binge drinking might be driven by inherent differences in gene expression between male and female mice brains, particularly within the CeA. Another limitation to our study is that we assay a limited number of genes, and did not assay protein changes.

To our knowledge, this is the first time that a study has tested what happens to a set of CeA genes during different epochs of abstinence, 1 day and 7 days, after a period of binge drinking in rodents of both sexes. The differences between male and female mice in binge drinking behavior and could be driven by differing motivation to consume alcohol, driven in part by engagement of differential signaling pathways in the CeA. In support of this, we have previously found that deletion of Oprk1 in the CeA has a greater impact on alcohol consumption in males as compared to female mice^22^. As the field moves forward with more studies balanced across sex of animal, we will be able to develop more detailed models of these potential sex differences. In keeping with this, our results strongly support the need to perform more direct mechanistic studies to better understand and target potential therapeutic or preventive pathways of alcohol use disorder in the CeA.

## Supplementary Figures

**Figure S1:**
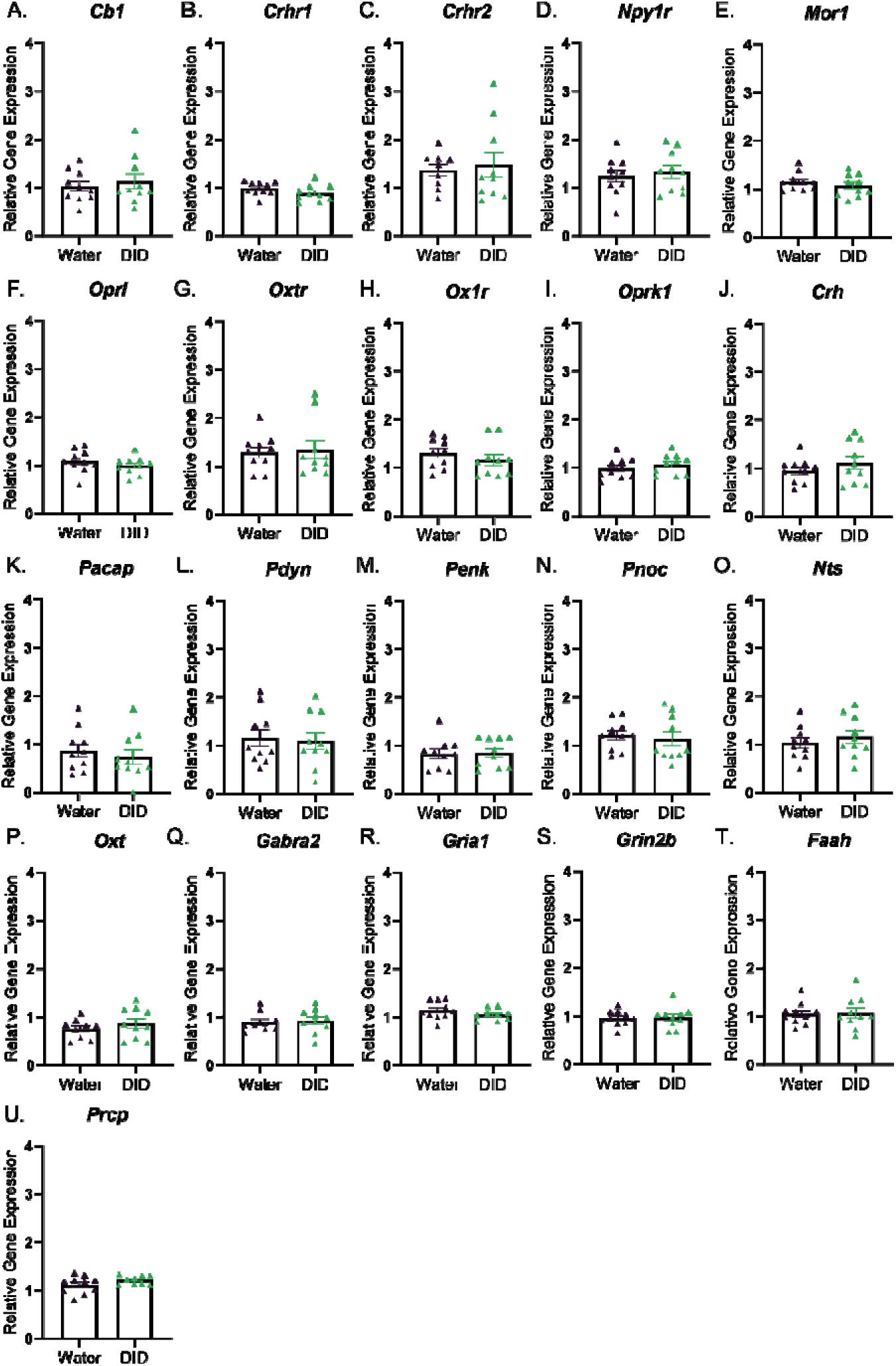
Relative gene expression to housekeeping gene actin B in acute abstinence after DID male mice. (**A-U**) Relative gene expression level of CeA derived genes from male mice from the acute abstinence group (1 day of abstinence after DID) compared to their respective water group. Error bars are depicted as ±SEM.

**Figure S2.**
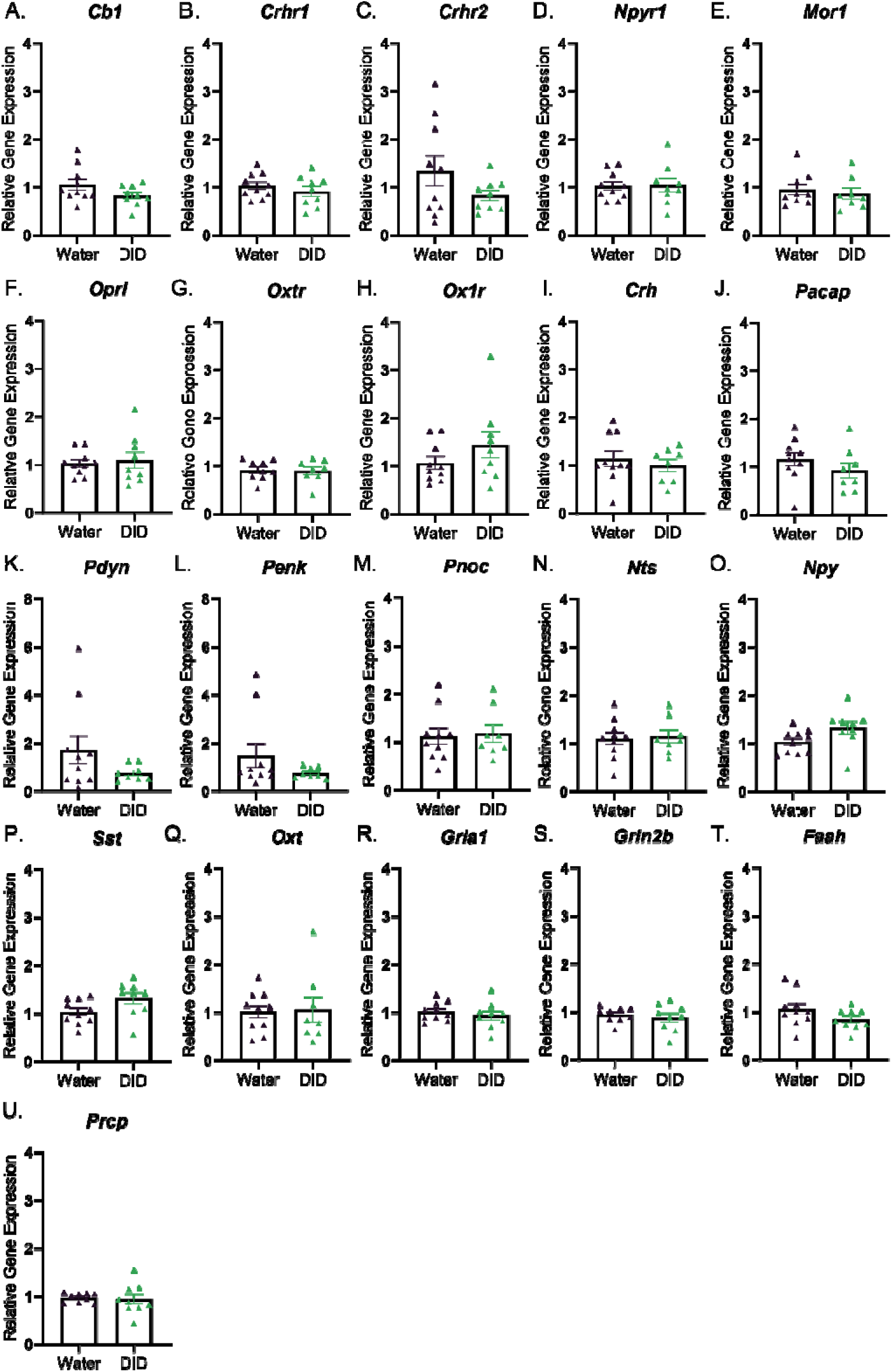
Relative gene expression to housekeeping gene actin B in protracted abstinence after DID male mice. (**A-U**) Relative gene expression level of CeA derived genes from male mice of the protracted abstinence group (7 days of abstinence after DID) compared to their respective water group. Error bars are depicted as ±SEM.

**Figure S3:**
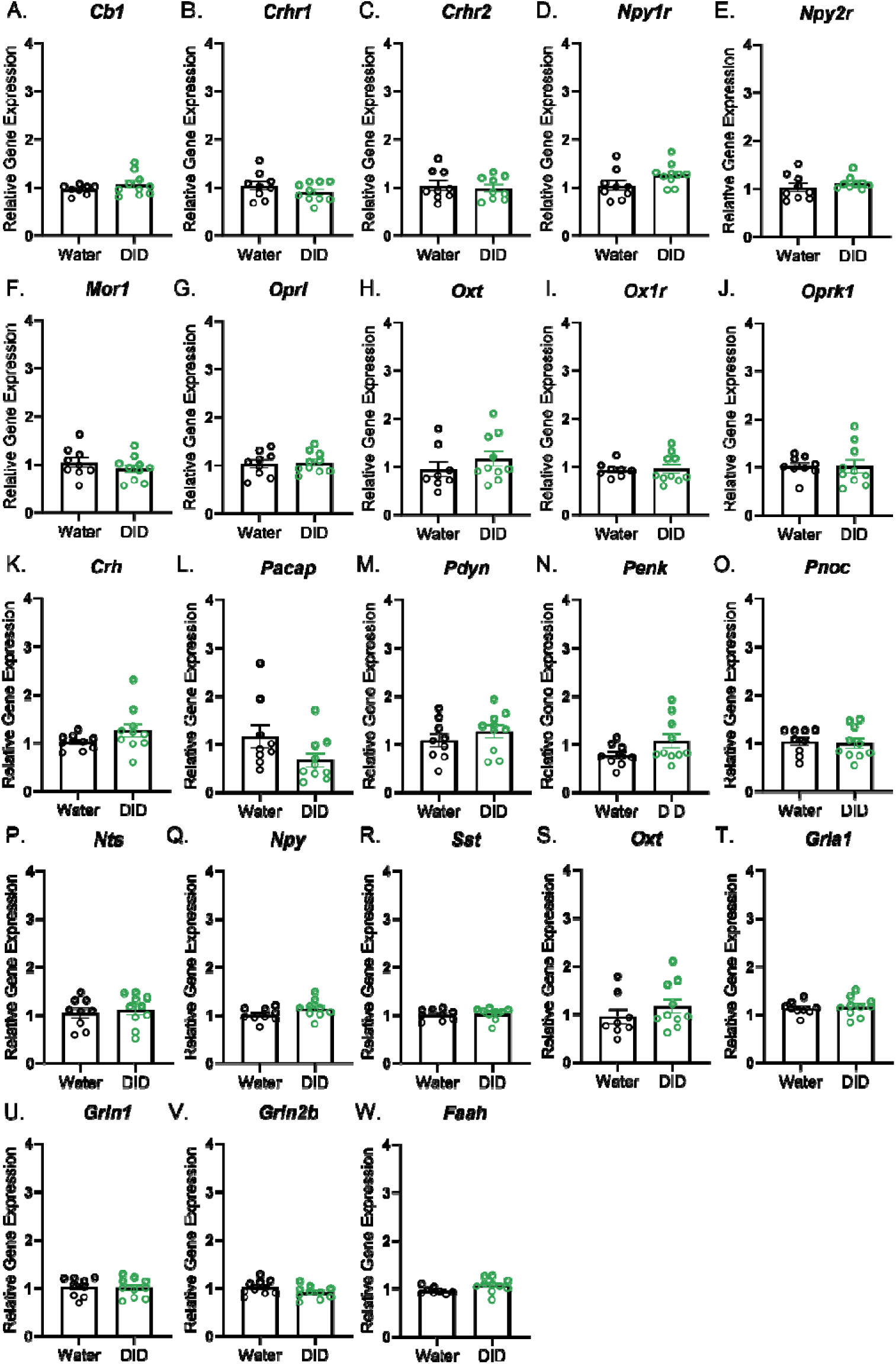
Relative gene expression to housekeeping gene actin B in acute abstinence after DID female mice. (**A-U**) Relative gene expression level of CeA derived genes from the female acute abstinence group (1 day of abstinence after DID) compared to their respective water group. Error bars are depicted as ±SEM.

**Figure S4:**
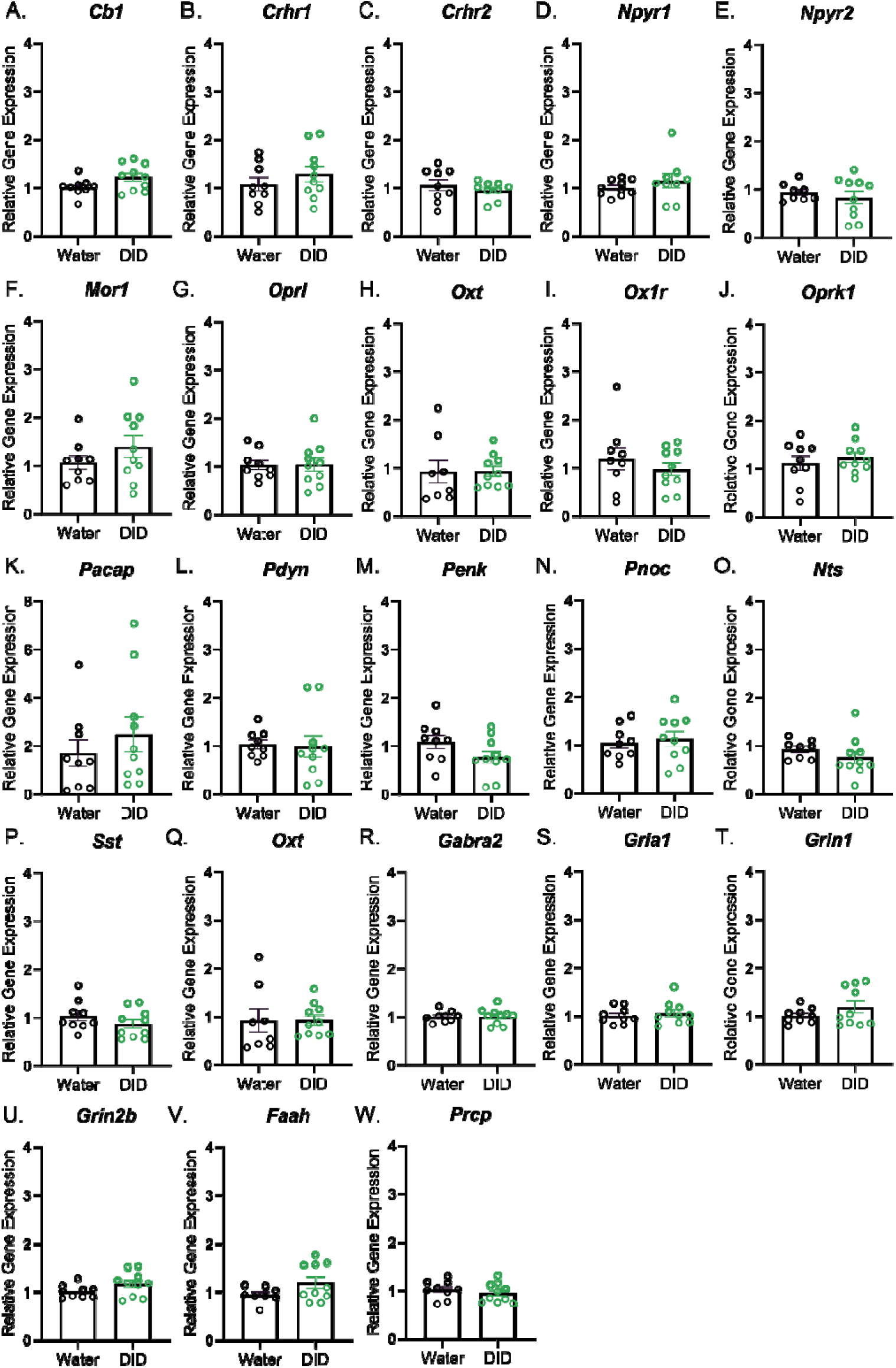
Relative gene expression to housekeeping gene actin B in protracted abstinence after DID female mice. (**A-U**) Relative gene expression level of CeA derived genes from the female protracted abstinence group (7 days of abstinence after DID) compared to their respective water group. Error bars are depicted as ±SEM.

## Acknowledgments

The authors thank Gabrielle Cannon at the UNC Advanced Analytics Core (Center for GI Biology and Disease; NIH funding number (P30 DK034987) for assistance with cDNA preamplification and qRT-PCR.

## References

1. Carvalho, A. F., Heilig, M., Perez, A., Probst, C. & Rehm, J. Alcohol use disorders. Lancet 394, 781–792 (2019).

2. Esser, M. B. et al. Estimated deaths attributable to excessive alcohol use among US adults aged 20 to 64 years, 2015 to 2019. JAMA Netw. Open 5, e2239485 (2022).

3. Sacks, J. J., Gonzales, K. R., Bouchery, E. E., Tomedi, L. E. & Brewer, R. D. 2010 national and state costs of excessive alcohol consumption. Am. J. Prev. Med. 49, e73– e79 (2015).

4. Benjamin, E. J. et al. Heart Disease and Stroke Statistics-2018 Update: A Report From the American Heart Association. Circulation 137, e67–e492 (2018).

5. Crossin, R. et al. The New Zealand drug harms ranking study: A multi-criteria decision analysis. J Psychopharmacol (Oxford) 37, 891–903 (2023).

6. Koob, G. F. & Volkow, N. D. Neurobiology of addiction: a neurocircuitry analysis. Lancet Psychiatry 3, 760–773 (2016).

7. Roberto, M., Kirson, D. & Khom, S. The role of the central amygdala in alcohol dependence. Cold Spring Harb. Perspect. Med. 11, (2021).

8. Greenfield, T. K. et al. Risks of alcohol use disorders related to drinking patterns in the U.S. general population. J. Stud. Alcohol Drugs 75, 319–327 (2014).

9. Lee, K. M., Coelho, M. A., Sern, K. R. & Szumlinski, K. K. Homer2 within the central nucleus of the amygdala modulates withdrawal-induced anxiety in a mouse model of binge-drinking. Neuropharmacology 128, 448–459 (2018).

10. Walker, L. C. A balancing act: the role of pro– and anti-stress peptides within the central amygdala in anxiety and alcohol use disorders. J. Neurochem. 157, 1615–1643 (2021).

11. Freeman, K. et al. Coordinated dynamic gene expression changes in the central nucleus of the amygdala during alcohol withdrawal. Alcohol. Clin. Exp. Res. 37 **Suppl 1**, E88–100 (2013).

12. McBride, W. J. et al. Changes in gene expression in regions of the extended amygdala of alcohol-preferring rats after binge-like alcohol drinking. Alcohol 44, 171–183 (2010).

13. Jesse, S. et al. Alcohol withdrawal syndrome: mechanisms, manifestations, and management. Acta Neurol. Scand. 135, 4–16 (2017).

14. Bloch, S. et al. Assessing negative affect in mice during abstinence from alcohol drinking: Limitations and future challenges. Alcohol 100, 41–56 (2022).

15. Koob, G. F. & Vendruscolo, L. Theoretical frameworks and mechanistic aspects of alcohol addiction: alcohol addiction as a reward deficit/stress surfeit disorder. Curr. Top. Behav. Neurosci. (2023) doi:10.1007/7854_2023_424.

16. Centanni, S. W., Bedse, G., Patel, S. & Winder, D. G. Driving the Downward Spiral: Alcohol-Induced Dysregulation of Extended Amygdala Circuits and Negative Affect. Alcohol. Clin. Exp. Res. 43, 2000–2013 (2019).

17. McBride, W. J. et al. Changes in gene expression within the extended amygdala following binge-like alcohol drinking by adolescent alcohol-preferring (P) rats. Pharmacol. Biochem. Behav. 117, 52–60 (2014).

18. Flanigan, M. E. et al. Subcortical serotonin 5HT2c receptor-containing neurons sex-specifically regulate binge-like alcohol consumption, social, and arousal behaviors in mice. Nat. Commun. 14, 1800 (2023).

19. Sparrow, A. M. et al. Central neuropeptide Y modulates binge-like ethanol drinking in C57BL/6J mice via Y1 and Y2 receptors. Neuropsychopharmacology 37, 1409–1421 (2012).

20. Robinson, S. L. & Thiele, T. E. Somatostatin signaling modulates binge drinking behavior via the central nucleus of the amygdala. Neuropharmacology 237, 109622 (2023).

21. Jin, Z. et al. Expression of specific ionotropic glutamate and GABA-A receptor subunits is decreased in central amygdala of alcoholics. Front. Cell. Neurosci. 8, 288 (2014).

22. Bloodgood, D. W. et al. Kappa opioid receptor and dynorphin signaling in the central amygdala regulates alcohol intake. Mol. Psychiatry 26, 2187–2199 (2021).

23. Anderson, R. I. et al. Dynorphin-kappa opioid receptor activity in the central amygdala modulates binge-like alcohol drinking in mice. Neuropsychopharmacology 44, 1084–1092 (2019).

24. Baird, M. A., Hsu, T. Y., Wang, R., Juarez, B. & Zweifel, L. S. κ Opioid Receptor-Dynorphin Signaling in the Central Amygdala Regulates Conditioned Threat Discrimination and Anxiety. eNeuro 8, (2021).

25. Vijay, A. et al. PET imaging reveals lower kappa opioid receptor availability in alcoholics but no effect of age. Neuropsychopharmacology 43, 2539–2547 (2018).

26. Kiefer, F. et al. Involvement of NMDA receptors in alcohol-mediated behavior: mice with reduced affinity of the NMDA R1 glycine binding site display an attenuated sensitivity to ethanol. Biol. Psychiatry 53, 345–351 (2003).

27. Xiang, Y., Kim, K.-Y., Gelernter, J., Park, I.-H. & Zhang, H. Ethanol upregulates NMDA receptor subunit gene expression in human embryonic stem cell-derived cortical neurons. PLoS ONE 10, e0134907 (2015).

28. Lieberman, R., Levine, E. S., Kranzler, H. R., Abreu, C. & Covault, J. Pilot study of iPS-derived neural cells to examine biologic effects of alcohol on human neurons in vitro. Alcohol. Clin. Exp. Res. 36, 1678–1687 (2012).

29. Ashton, M. K. et al. Sex differences in GABAA receptor subunit transcript expression are mediated by genotype in subjects with alcohol-related cirrhosis of the liver. Genes Brain Behav. 21, e12785 (2022).

30. Liu, J. et al. Binge alcohol drinking is associated with GABAA alpha2-regulated Toll-like receptor 4 (TLR4) expression in the central amygdala. Proc Natl Acad Sci USA 108, 4465–4470 (2011).

31. Bruschetta, G., Jin, S., Kim, J. D. & Diano, S. Prolyl carboxypeptidase in Agouti-related Peptide neurons modulates food intake and body weight. Mol. Metab. 10, 28–38 (2018).

32. Diano, S. New aspects of melanocortin signaling: a role for PRCP in α-MSH degradation. Front. Neuroendocrinol. 32, 70–83 (2011).

33. Lerma-Cabrera, J. M. et al. Adolescent binge-like ethanol exposure reduces basal α-MSH expression in the hypothalamus and the amygdala of adult rats. Pharmacol. Biochem. Behav. 110, 66–74 (2013).

